# The Phenoscape Knowledgebase: tools and APIs for computing across phenotypes from evolutionary diversity and model organisms

**DOI:** 10.1101/071951

**Authors:** James P. Balhoff

## I. Introduction

The Phenoscape Knowledgebase (KB) is an ontology-driven database that combines existing phenotype annotations from model organism databases with new phenotype annotations from the evolutionary literature. Phenoscape curators have created phenotype annotations for more than 5,000 species and higher taxa, by defining computable phenotype concepts for more than 20,000 character states from over 160 published phylogenetic studies. These phenotype concepts are in the form of Entity–Quality (EQ) [1] compositions which incorporate terms from the Uberon anatomy ontology, the Biospatial Ontology (BSPO), and the Phenotype and Trait Ontology (PATO). Taxonomic concepts are drawn from the Vertebrate Taxonomy Ontology (VTO). This knowledge of comparative biodiversity is linked to potentially relevant developmental genetic mechanisms by importing associations of genes to phenotypic effects and gene expression locations from zebrafish (ZFIN [2]), mouse (MGI [3]), Xenopus (Xenbase [4]), and human (Human Phenotype Ontology project [5]). Thus far, the Phenoscape KB has been used to identify candidate genes for evolutionary phenotypes [6], to match profiles of ancestral evolutionary variation with gene phenotype profiles [7], and to combine data across many evolutionary studies by inferring indirectly asserted values within synthetic supermatrices [8]. Here we describe the software architecture of the Phenoscape KB, including data ingestion, integration of OWL reasoning, web service interface, and application features (Fig. 1).

**Fig. 1.**
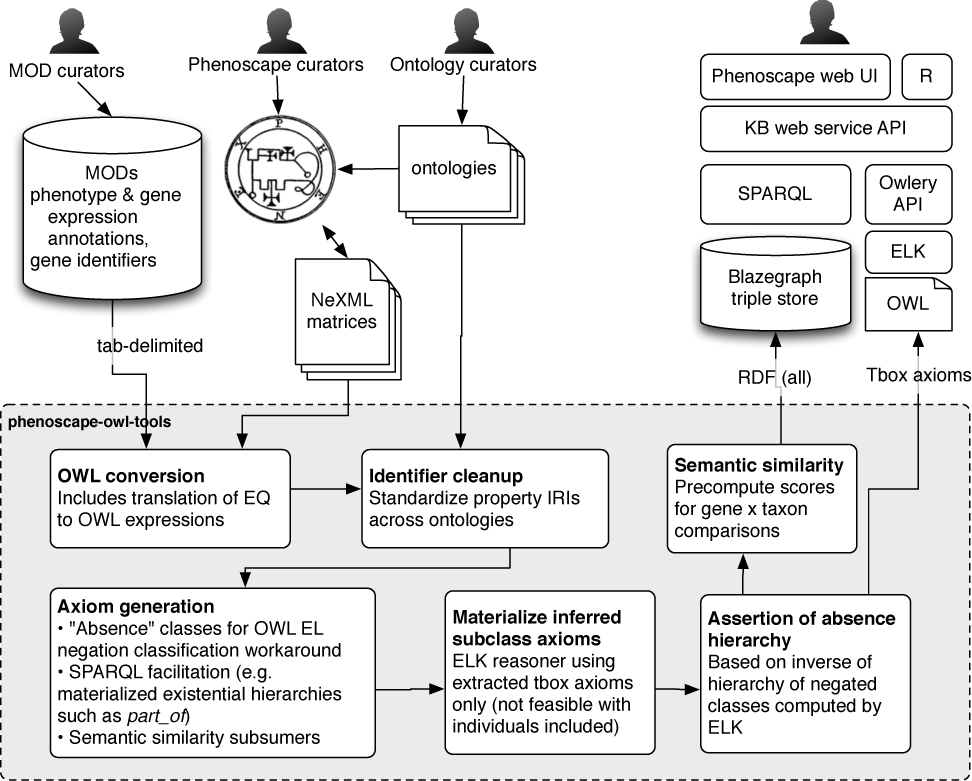
Phenoscape KB software architecture and data flow.

## II. Data ingestion and inference materialization

Phenoscape curators annotate character by taxon matrices associated with phylogenetic publications using the Phenex software [9], following the curation process described by Dahdul et al. [10]. The resulting NeXML data files, along with tab-delimited data dumps obtained from model organism databases, are translated into OWL models (including definitions of EQ phenotypes) using the Phenoscape data ingest pipeline (phenoscape-owl-tools, available with all other described software under an MIT open source license in the Phenoscape GitHub repository, https://github.com/phenoscape). OWL transformation code for individual data sources is kept concise and readable by using a Scala-based domain specific language for OWL axioms (Scowl) [11]. Next, all OWL data files, along with referenced ontologies, are processed to standardize property IRIs; in many cases community ontologies use non-standard or competing IRIs for the “same” property, e.g. our system accounts for ten variants of the *part_of* property. We rename these, rather than assert equivalence axioms, to simplify downstream reasoning and querying.

Based on the content of the data and reference ontologies, several “ontologies” are programmatically generated, for the purpose of precomputing inferred concept hierarchies supporting various Phenoscape use cases: 1) materialization of the transitive closure for selected properties; 2) generation of grouping phenotypes for semantic similarity queries; 3) generation of “absence” concepts for custom negation reasoning. The ELK reasoner [12] is then used to compute the inferred classification hierarchy, and inferred subclass axioms are materialized into concrete assertions. Because ELK does not support negation reasoning, we have also implemented a custom procedure to compute a class hierarchy for a predetermined set of negations [13]. To make reasoning on this large dataset feasible, we extract only the class axioms (Tbox) for input into ELK. This is sufficient for our purposes since most of the data is in the form of compositional class expressions; however it does restrict use cases that would rely on inference of property assertions or instance classification. All data, including asserted and inferred class axioms as well as instance data, are loaded into the Blazegraph RDF triple store [14], constituting approximately 100 million RDF triples. A separate OWL file including only class axioms (~870,000 logical axioms) is saved for later use in reasoner queries.

## III. Semantic similarity precomputation

Phenotypic profiles for evolutionarily variable taxonomic nodes and model organism genes are computed as described in [7], and loaded into Blazegraph. The inferred phenotype class hierarchy is used to precompute semantic similarity scores between all pairs of evolutionary variation profiles and gene phenotype profiles, using an information content-based metric. The set of pair comparisons is broken into chunks and processed in parallel on a compute cluster to reduce computation time. The resulting set of similarity scores, along with computed statistical support, is loaded into Blazegraph, resulting in a final triple store totaling approximately 300 million RDF triples.

## IV. Web service interfaces

Access to data in the Phenoscape KB is provided by two web service applications, implemented in Scala using the Spray HTTP toolkit [15]. The first, Owlery, provides a generic JSON-format API to an any OWL API-based reasoner; for the Phenoscape KB we use ELK, loaded with extracted class axioms from the ontologies and data. We use Owlery to support web application queries that require reasoning on arbitrary OWL class expressions. Owlet provides web services for description logic queries (obtaining subclasses and superclasses), and also supports reasoner-based query expansion using our Owlet package. The Owlery API is documented at http://docs.owlery.apiary.io/. The second web service application, phenoscape-kb-services, is the primary public Phenoscape API and provides Phenoscape application-specific services such as annotation query, semantic similarity, annotation support for presence/absence inference, and term info. Phenoscape-kb-services obtains most results via SPARQL query to the Blazegraph triple store. For use cases requiring computation by the ELK reasoner, SPARQL queries with embedded OWL expressions are first expanded using the Owlet service provided by the Owlery API before being submitted to Blazegraph. The phenoscape-kb-services API returns most results in both JSON and tab-delimited text formats. Documentation for the phenoscape-kb-services API can be found at http://docs.phenoscapekb.apiary.io/.

## V. Applications

A public web user interface for the Phenoscape Knowledgebase is available at http://kb.phenoscape.org/. The web interface is a client-side browser application developed in the AngularJS JavaScript framework [16]. The application is implemented entirely upon web service calls to the phenoscape-kb-services API, ensuring that all functionality is available via a documented, public API. Using the web application, researchers can query evolutionary descriptions and gene phenotypes relevant to particular structures and qualities, search for taxonomic groups exhibiting variation similar to the phenotypic effects of a gene of interest [7] (and vice versa), and export synthetic presence/absence supermatrices using the OntoTrace system [8]. Building upon the same web service API, we have also implemented an R package, rPhenoscape, which makes some of the functionality of the KB available within the R statistical computing environment.

## VI. Conclusion

The Phenoscape Knowledgebase architecture illustrates one approach to integration of multiple datasets in a rich ontological framework. Deriving the full benefit of the sophisticated knowledge represented in OBO library ontologies will often require application of automated reasoners and programmatic generation of axioms and concepts to facilitate particular use cases. Here we have provided an overview of the mix of reusable components and special purpose code we have developed to support Phenoscape; it is our hope that continued evolution of the Phenoscape KB architecture will result in further identification and development of reusable tools which can support similar efforts.

## Footnotes

Funded by National Science Foundation grants DBI-1062404, DBI-1062542.

